# Determination of growth-coupling strategies and their underlying principles

**DOI:** 10.1101/258996

**Authors:** Tobias B. Alter, Lars M. Blank, Birgitta E. Ebert

## Abstract

Metabolic coupling of product synthesis and microbial growth is a prominent approach for maximizing production performance. Growth-coupling (GC) also helps stabilizing target production and allows the selection of superior production strains by adaptive laboratory evolution. We have developed the computational tool gcOpt, which identifies knockout strategies leading to the best possible GC by maximizing the minimally guaranteed product yield. gcOpt implicitly favors solutions resulting in strict coupling of product synthesis to growth and metabolic activity while avoiding solutions inferring weak, conditional coupling.

GC intervention strategies identified by gcOpt were examined for GC generating principles under diverse conditions. Curtailing the metabolism to render product formation an essential carbon drain was identified as one major strategy generating strong coupling of metabolic activity and target synthesis. Impeding the balancing of cofactors and protons in the absence of target production was the underlying principle of all other strategies and further increased the GC strength of the aforementioned strategies. Thus, generating a dependency between supply of global metabolic cofactors and product synthesis appears to be advantageous in enforcing strong GC.

**Abbreviations:**

ATP: Adenosine triphosphate
ATPM: ATP maintenance requirements reaction
ATPcsc: Carbon specific ATP synthesis capability
ATPsc: ATP synthesis capability
CoA: Coenzyme A
EM: Elementary mode
EMA: Elementary modes analysis
FBA: Flux balance analysis
GC: Growth-coupling
GCS: Growth-coupling strength
H^+^: Proton
hGC: Holistic growth-coupling
MCS: Minimal cut sets
MILP: Mixed integer linear program
NAD^+^: Nicotinamide adenine dinucleotide (oxidized)
NADH: Nicotinamide adenine dinucleotide (reduced)
NADP^+^: Nicotinamide adenine dinucleotide phosphate (oxidized)
NADPH: Nicotinamide adenine dinucleotide phosphate (reduced)
NGAM: Non-growth associated maintenance
Pi: Phosphate molecule
PPP: Pentose phosphate pathway
sGC: Strong growth-coupling
wGC: Weak growth-coupling

## Introduction

Metabolic engineering approaches strive to optimize microbial cell-factories for robust, profitable, and sustainable industrial applications (Nakamura & Whited, 2003). One applied principle within this field of research is to metabolically couple the synthesis of the product of interest to microbial growth by appropriate genetic modifications (Fong *et al,* 2005; Jantama *et al,* 2008; Trinh *et al,* 2008; Jiang *et al,* 2013; Layton & Trinh, 2014). The main motivation in generating growth-coupled production is to shift the tug of war for the substrate carbon towards the synthesis of the desired chemical (Kashket & Zhi-Yi Cao, 1995; Van Dien, 2013; Jouhten *et al,* 2017). Consequently, growth-coupling (GC) efficiently facilitates the use of well-established adaptive laboratory evolution methods for production strain optimization purposes by employing growth as a simple selection criterion (Portnoy *et al,* 2011; Sandberg *et al,* 2017).

Three distinct GC phenotypes differing in GC strength can be distinguished, which become apparent from computing and plotting so-called metabolic yield spaces (Feist *et al,* 2010). These yield spaces are projections of the accessible flux space onto the 2D plane spanned by the growth rate and the yield of the target product on the main carbon substrate (Fig. 1). The lower limit of a yield space depicts the minimally guaranteed product yield for the accessible range of growth rates. Hence, a lower bound greater than zero for a particular growth state directly implies GC. In the following, yield spaces, in which a product yield greater zero only occurs at elevated growth rates, will be denoted as a weak GC (wGC) characteristic (Fig. 1 A). For *Saccharomyces cerevisiae* and *Escherichia coli*, for example, such a wGC is naturally observed for fermentation products, *e.g.*, ethanol or acetate, under anaerobic conditions or during overflow metabolism. By this means, holistic GC (hGC) is encountered if the lower yield bound is above zero for all growth rates greater than zero (Fig. 1 B) while strong GC (sGC) is referred to yield spaces showing a mandatorily active target compound production for *all* metabolic states including zero growth (Fig. 1 C).

**Figure 1.**
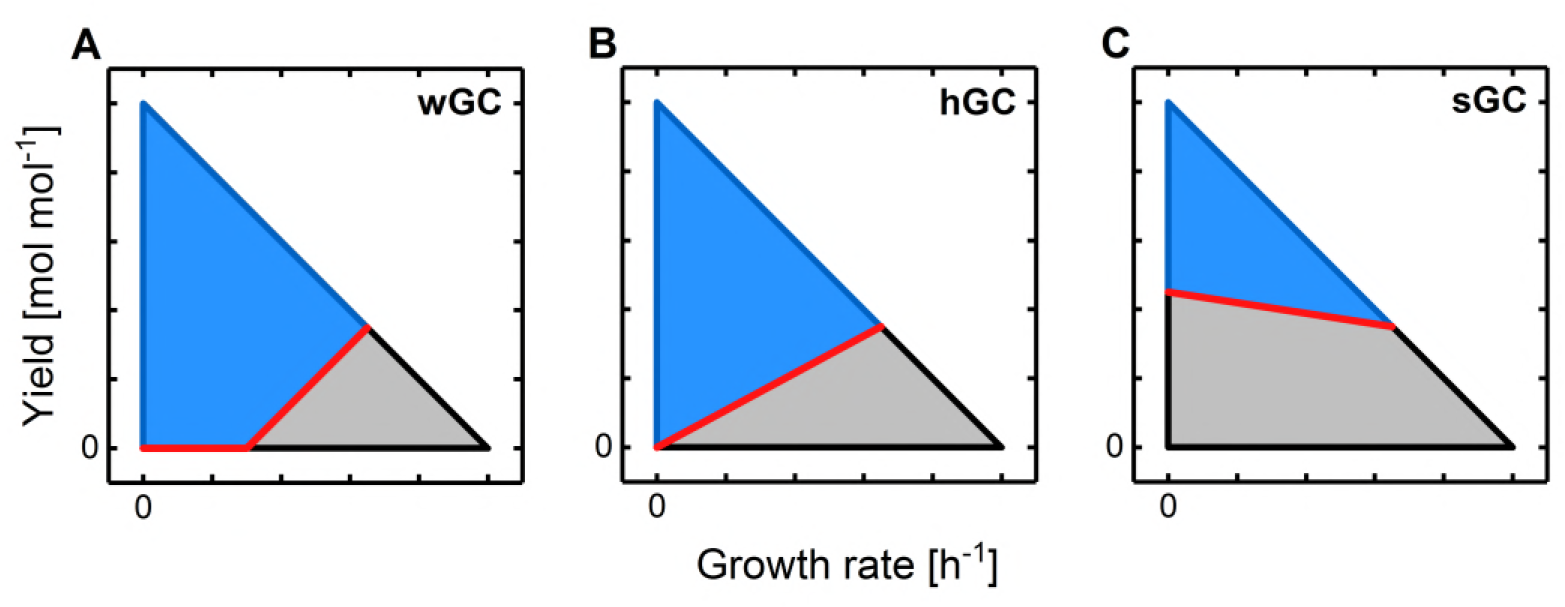
Exemplary yield spaces showing three distinct types of GC. A Projection of the accessible flux space onto the 2D plane spanned by the growth rate and the yield of a target product on the main carbon substrate after applying respective reaction deletions (blue area). The grey area represents the yield space of the wild-type strain which is inaccessible for the mutant strain. The lower yield bound, hence the minimally guaranteed product yield, is marked by the red line. Here, a weak GC (wGC) characteristic is shown since product yield is only guaranteed for elevated growth rates. B Yield space representing holistic GC (hGC). Target compound production is mandatorily active for all growth states but zero growth. C Here, strong GC (sGC) is shown. The lower yield bound is strictly greater zero and thus, for all metabolic states target compound production is enforced.

Various computational algorithms exploiting the rich information content of stoichiometric metabolic models have been developed to specifically provide reaction deletion strategies leading to GC. These approaches are generally grouped into Flux Balance Analysis (FBA) and Elementary Modes Analysis (EMA) based methods. Classical FBA focuses on a particular metabolic phenotype by optimizing a biological meaningful objective function subject to steady-state mass balance constraints (Savinell & Palsson, 1992). Thus, GC strain designs identified by FBA-based frameworks such as OptKnock (Burgard *et al*, 2003) and RobustKnock (Tepper & Shlomi, 2009) enforce GC at only distinct metabolic states, which is maximal biomass formation in the given examples. While OptKnock attempts to solely maximize the target compound production, RobustKnock maximizes a *minimally guaranteed* production and thus enforces GC at maximal growth. Complementarily to FBA, EMA utilizes the nondecomposable steady-state flux distributions, called elementary modes (EMs), of a metabolic network from which any feasible flux state can be derived by linear combinations of these EMs (Schuster *et al,* 1999; Gagneur & Klamt, 2004). By exploiting the nondecomposibility feature of EMs, minimal sets of reaction deletions, coined minimal cut sets (MCSs), can be identified that disable all EMs responsible for undesired metabolic functionalities (Klamt & Gilles, 2004). Methods such as constrained minimal cut sets (Hädicke & Klamt, 2011; von Kamp & Klamt, 2017) or minimal metabolic functionality (Trinh *et al,* 2008) use EMA to determine MCSs, which remove all EMs producing only biomass and hence lead to GC. The main disadvantage of EMA-based compared to FBA-based methods is the computationally expensive necessity to enumerate all EMs, thus limiting the application of EMA to small or mid-scale metabolic networks. Recently, this has been overcome by introducing MCSEnumerator, an algorithm that sequentially enumerates the smallest MCSs and significantly reduces the computational costs (von Kamp & Klamt, 2014). Since the underlying constrained MCS method requires the definition of a minimal bound on growth rate and target product yield, MCSEnumerator specifically searches for intervention strategies leading to sGC (Fig. 1 C). This, however, may result in neglection of the best possible but suboptimal solutions resulting in wGC or hGC when no sGC solutions exist for the user-defined maximum allowable number of reaction deletions. To effectively gain from the advantages of different methods in terms of a biologically robust strain design, combinations and adaptions of the mentioned algorithms have been reported (Feist *et al,* 2010; Shabestary & Hudson, 2016; Nair *et al,* 2017).

Beside the *in silico* identification of GC intervention strategies, research on the general feasibility and driving forces of the coupling between growth and target product synthesis has been conducted. Based on the EMA approach, Klamt & Mahadevan (2015) have built a theoretical framework to relate GC to the existence of elementary modes and vectors that fulfill specific requirements on biomass and product yields. By applying this framework to a metabolic model of the central carbon metabolism of *E. coli* (Trinh *et al,* 2008; Hädicke & Klamt, 2011), they were able to show that synthesis of any metabolite can be coupled to growth. Recently, Jouhten *et al.* (2017) proposed a biochemical basis for the generation of growth-coupled product synthesis. They introduced the concept of anchor reactions, which split the substrate carbon among one or more biomass precursors and the target compound. Existence of an anchor reaction that is or can be made essential for the synthesis of a biomass precursor thus implies feasibility of growth-product coupling. This has similarly been expressed by Klamt and Mahadevan (2015) in the requirement for at least one elementary mode allowing for both growth and product synthesis. In contrast, it was claimed elsewhere that GC results from an induced imbalance of reduction or energy equivalents, which can only be overcome by active product synthesis (Erdrich *et al,* 2014; Shabestary & Hudson, 2016; Jiang *et al,* 2013). Erdrich *et al.* (2014) pointed out, that this imbalance is particularly pronounced under anaerobic conditions where oxygen as final electron acceptor is missing and ATP generation is naturally limited mainly to fermentation pathways and glycolysis.

In view of these disparate explanations for GC, we aimed at further unraveling key principles of reaction deletion strategies leading to GC by identifying relevant genetic intervention strategies for a set of metabolites and investigating the specific operating principle of these strategies. We developed the FBA-based algorithm gcOpt, which determines the best possible GC knockout strategy for a given target compound, a specific substrate and a defined maximum number of reaction deletions. gcOpt was applied to calculate GC intervention strategies for a broad range of metabolites of a core as well as a genome-scale metabolic model of *E. coli.* The resulting strategies were subsequently examined regarding the consequence of imposed growth-coupled product synthesis on metabolic network operation.

## Results

### gcOpt prioritizes strain designs resulting in strong growth coupling

The pursued approach to identify GC strain designs with maximal possible GC strength was derived from the yield space representation of GC mutants (cf. Fig 1). While the GC classification into wGC, hGC, and sGC provides a qualitative notion of the GC strength, the position of the lower yield boundary can be interpreted as a quantitative measure: the higher the boundary in terms of positive yield values, the stronger the GC. The shape of this yield space boundary along the growth rate axis is not arbitrary. It is rather a part of the hull giving the admissible flux space and, since the flux space is determined by a linear equation system, the lower yield space boundary is convex (Schuster & Hilgetag, 1994). It follows from the convexity property that by increasing the lower yield for one specific growth rate by, *e.g.*, deletion of one or more reactions, the lower yield boundary at all admissible growth rates is raised, resulting in an overall increase of the GC strength. This principle was implemented in a bi-level optimization algorithm, gcOpt, which maximizes the minimum product yield of a target compound at a fixed, medium growth rate using appropriate reaction deletions (Fig 2). Ultimately, gcOpt provides strain designs with the best possible GC for a given metabolic network and the defined maximum number of modifications (see the Methods section for a detailed description and formulation of gcOpt).

**Figure 2.**
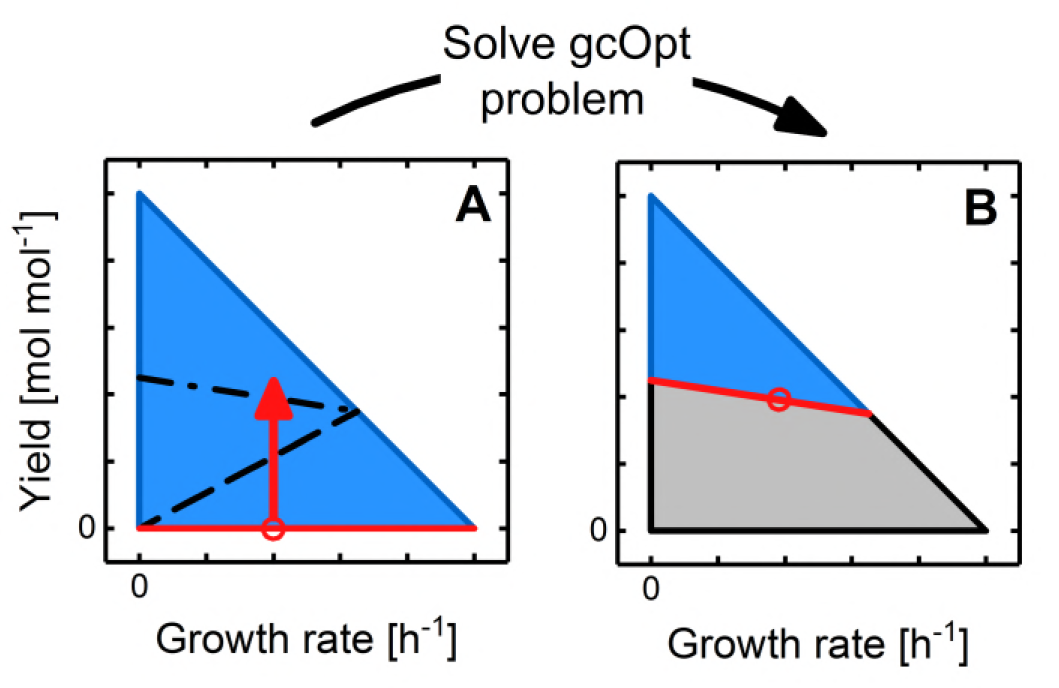
Schematic principle of gcOpt. A Exemplary yield space of a wild-type strain showing no GC. The blue area denotes the admissible flux space as a function of the yield and growth rate. The red, horizontal line marks the lower yield bound, whereas the black dashed lines show the lower yield bounds of, possible mutant strains with differing GC characteristics. The red arrow denotes the optimization principle of gcOpt, which is maximization of the minimally guaranteed yield at a medium fixed growth rate. B Yield space of a reaction deletion mutant strain showing the best possible GC. In addition to **A**, the grey area depicts the flux space that is inadmissible to the mutant strain derived the deletion strategy identified by gcOpt.

Identification of strain designs leading to GC of ethanol production in *E. coli* under anaerobic conditions was used to demonstrate the functionality of gcOpt. This classic example has already been investigated by applying diverse computational methods (Trinh *et al,* 2008, Hädicke & Klamt, 2011) to a metabolic model of the central carbon metabolism of *E. coli*, here referred to as CT86 (Trinh *et al,* 2008). Using CT86, gcOpt was applied allowing maximum numbers of reaction deletions from one to five at three different fixed growth rates *µ_fix_* of 0.01 *h*^−1^, 0.1 *h*^−1^ and 0.25 *h*^−1^. Anaerobic growth on glucose was simulated by setting the maximum glucose and oxygen uptake rate to 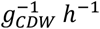 and zero, respectively. The designs identified by gcOpt (Fig. 3 A-C) clearly indicate that the lower yield bound, and hence the GC strength, increased with a growing number of simultaneous reaction deletions while the maximal growth rate decreased and approached the chosen *µ_fix_.* The most extremely trimmed yield space was computed for the triple, quadruple and quintuple mutants at a *µ_fix_* of 0.01 *h*^−1^ (Fig. 3 A). The maximum growth rates did not exceed values of 0.05 *h*^−1^ while an ethanol yield of 1.7 *mol mol*^−1^ was strictly guaranteed implicating a tight metabolic coupling of growth and ethanol production. The quintuple mutant design computed for *µ_fix_ =* 0.25 *h*^−1^ (Fig. 3 C) was interesting in that it enforced a high ethanol yield at a relatively high maximal growth rate of 0.31 *h*^−1^. The minimally guaranteed yield was 0.85 *mol mol*^−1^, thus pointing to an excellent combination of GC and viability of this mutant. The predicted intervention strategies at a *µ_fix_ =* 0.1 *h*^−1^ (Fig. 3 B) were a good compromise between this and the extremely trimmed strain designs at a *µ_fix_ =* 0.01 *h*^−1^ with guaranteed yields of approx. *12 mol mol*^−1^ and maximal growth rates of 0.13 *h*^−1^. Fig. 3 D opposes yield spaces of GC strain designs found by various other methods to those identified by gcOpt (Fig. 3 A-C). By consulting the lower bounds of the yield spaces as a measure for the GC strength, the double and quadruple mutants determined by OptKnock, RobustKnock and cMCS, respectively, generally showed inferior GC characteristics than mutants of the same intervention sizes found by gcOpt. Moreover, although cMCS and MMF identified a tight GC for the quintuple and septuple mutants, for both mutant strains the product yield at maximal growth could take a range of values. A bottleneck in biomass precursor supply at elevated growth rates can be assumed in these cases since such edges of flux polyhedra in general, and thus of yield spaces in particular, correspond to flux capacity constraints (Klamt & Mahadevan, 2015). Such a phenomenon, however, was not seen for any gcOpt strain design and thus might be avoided by this algorithm.

**Figure 3.**
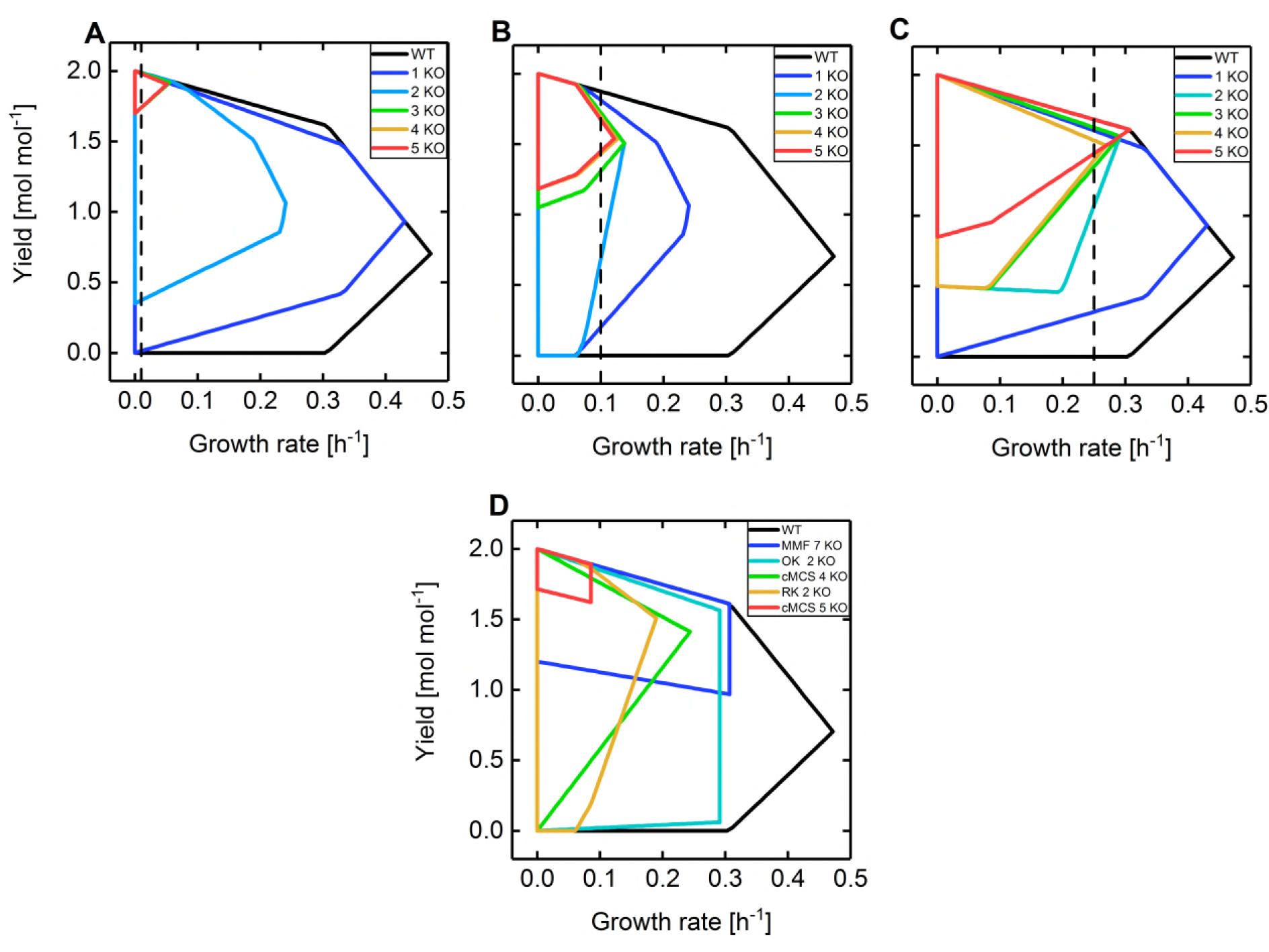
Ethanol yield spaces of GC strain designs identified by gcOpt (A-C) and several methods reported in the literature (D). A-C Ethanol yield spaces of the wild-type (black lines) and single to quintuple deletion mutant strains (colored lines) computed by gcOpt using the CT86 metabolic model. The vertical black dashed lines mark the chosen fixed growth rates *µ_fix_* for the respective computations (0.01 *h*^−1^ (**A**), 0.1 *h*^−1^ (**B**) and 0.25 *h*^−1^ (**C**)). D Ethanol yield spaces of the wild-type and several strain designs reported in the literature. The intervention strategies were identified using different computational approaches including MMF (Trinh *et al,* 2008) as well as OptKnock (OK), RobustKnock (RK), and constrained minimal cut sets (cMCS) (Hädicke & Klamt, 2011).

Consequently, gcOpt offers the advantage to compute the *best possible* GC strain design for a given microbial host, target compound, environmental condition and specified maximum number of genetic interventions. Moreover, the inherent approach of increasing the minimum product yield enforces the generally preferred sGC and hGC solutions, which guarantee product synthesis with growing or metabolically active organisms. This is a beneficial trait compared to alternative FBA based algorithms such as OptKnock or RobustKnock, which *per se* do not favor these designs over wGC solutions.

### Are there metabolic principles leading to growth-coupling?

Next, we applied gcOpt to compute a comprehensive dataset of GC designs, which we analyzed in-depth to decipher general metabolic principles that trigger GC. To this end, we computed intervention strategies with one to seven reaction deletions for the 36 central carbon metabolites of the *E. coli* iAF1260 core model under aerobic as well as anaerobic conditions. For both conditions, gcOpt simulations were additionally conducted with a decreased as well as an increased non-growth associated maintenance (NGAM) ATP requirement by changing the lower flux bound of the corresponding ATP maintenance requirement reaction (ATPM, Eq. 1) about 50 % from its standard value of 8.39 *mmol g*^−1^*h*^−1^ (Feist *et al,* 2007) to 4.2 *mmol g*^−1^*h*^−1^ and 12.2 *mmol g*^−1^*h*^−1^, respectively.

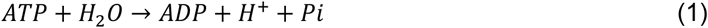

Equivalently to simulating the influence of the NGAM demand on finding GC strain designs, NGAM reactions were separately introduced for NAD(^+^/H) and NADP(^+^/H), virtually resembling an elevated turnover of these cofactors (Eq. 2–3). The fluxes were arbitrarily fixed to 5.0 *mmol g*^−1^*h*^−1^ or -5.0 *mmol g*^−1^*h*^−1^ to simulate the consumption of the oxidized or reduced cofactor, respectively.

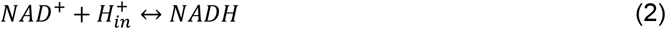

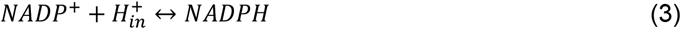

Equations 2 and 3 are mass but not charge balanced. It is assumed that the electrons are transferred from/to an imaginary electron donor/acceptor, which can be freely reduced or oxidized to avoid mass imbalances of additionally included redox cofactors. For each altered ATPM and virtual cofactor NGAM, GC strain designs were successfully identified for ca. 90 % of all metabolites under aerobic conditions, except for the condition of increased NADP^+^ consumption, which reduced the couplable metabolites to 75 % (Table EV1). The four metabolites, which could not be coupled to growth, were acetyl-CoA and succinyl-CoA, due to the model’s inability to compensate for the CoA drain, acetyl phosphate, and L-glutamine. For anaerobic growth, the percentage of growth-coupled metabolites was much lower (Table EV1). Interestingly, metabolites, for which gcOpt computed only wGC designs for standard conditions, could be strongly growth-coupled when the ATP NGAM was reduced. Among those were, *e.g.,* phosphorylated intermediates of glycolysis such as glucose-6-phosphate and 2-phosphoglycerate. The growth-coupled synthesis of those metabolites was apparently fueled by excess ATP.

To more quantitatively compare GC characteristics between different designs, a novel measure for the GC strength, termed as GCS, was introduced (cf. Methods). GCS relates the area of the accessible yield space of the wild-type strain to the inaccessible or blocked area below the lower yield bound of the mutant strain up to the maximum growth rate of the mutant (cf. Fig. 4). Thus, the higher the lower yield boundary of the mutant, the higher the GCS. The minimally guaranteed production rate of the target compound at maximal growth of the mutant strain is considered as an additional factor for determining the GCS (Eq. 6). Fig. 5 shows the mean GCS of all investigated metabolites for an increasing number of maximal reaction deletions for anaerobic as well as aerobic conditions and for altered or additionally introduced cofactor requirements. If no GC strain design was identified for a metabolite, the GCS was set to -2, defined as a complete lack of a coupling between growth and product synthesis. Under anaerobic conditions (Fig. 5 A), the mean GCS steadily increased with cumulative reaction knockouts from one to four and reached a plateau above this threshold for all investigated conditions. As already apparent form the increased number of sGC designs (Tab. 1), a reduced ATP demand, *i.e*., a low NGAM requirement, increased the mean GCS while alterations of the demand of the redox cofactors NAD(P/H) did not have a comparable effect. For aerobic conditions, we found coupling strategies with significantly higher mean GCS values. Both the number of metabolites and GCS of identified strategies were rather indifferent to alterations of the energy and redox cofactor demand (Fig. 5 B). Again, the mean GCS steadily increased with a growing number of reaction knockouts. The increase attenuated but did not reach a plateau in simulations restricted to maximal seven reaction deletions.

**Figure 4.**
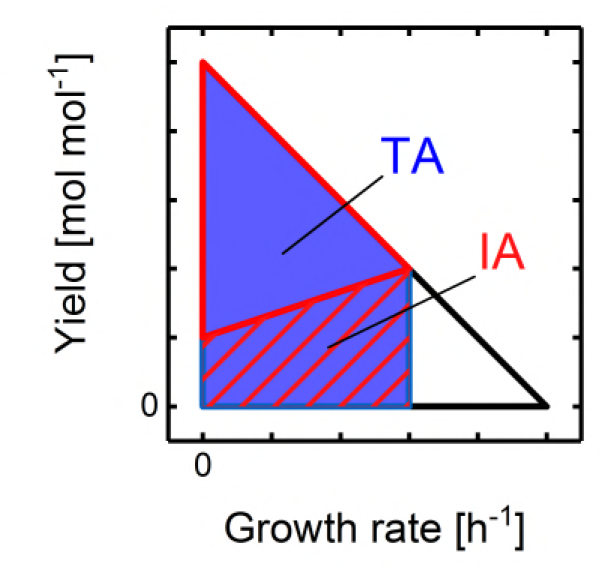
Illustration of the yield space areas used for calculating GCS. Scheme of a wild-type yield space showing no GC (black hull curve) and a GC strain design (red hull curve). The blue area TA illustrates the yield space of the wild-type up to the maximal growth rate of the mutant strain. The inaccessible yield space IA below the lower yield bound of the mutant is marked by the red hatched area.

**Figure 5.**
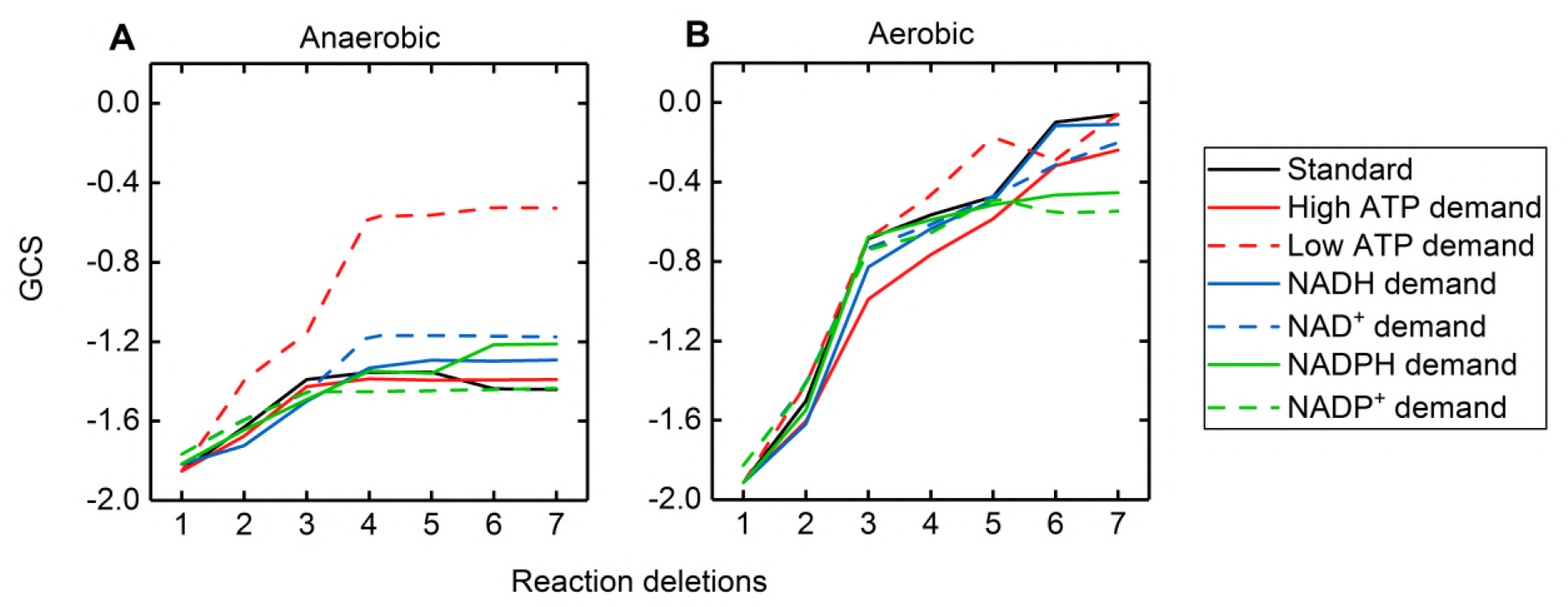
Mean GCS progressions as a function of the number of reaction deletions. A Mean GCS for increasing numbers of simultaneous reaction deletions of all GC strain designs identified by gcOpt for all metabolites of the *E. coli i*AF1260 core model under anaerobic conditions. The different lines embrace independent simulations applying a particular cofactor demand as illustrated by the legend. For the standard condition (black line) a NGAM ATP requirement of 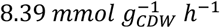 and no NADH, NAD, NADPH or NADP demand reaction was considered. B Cf. A, but for aerobic conditions.

**Table 1.**
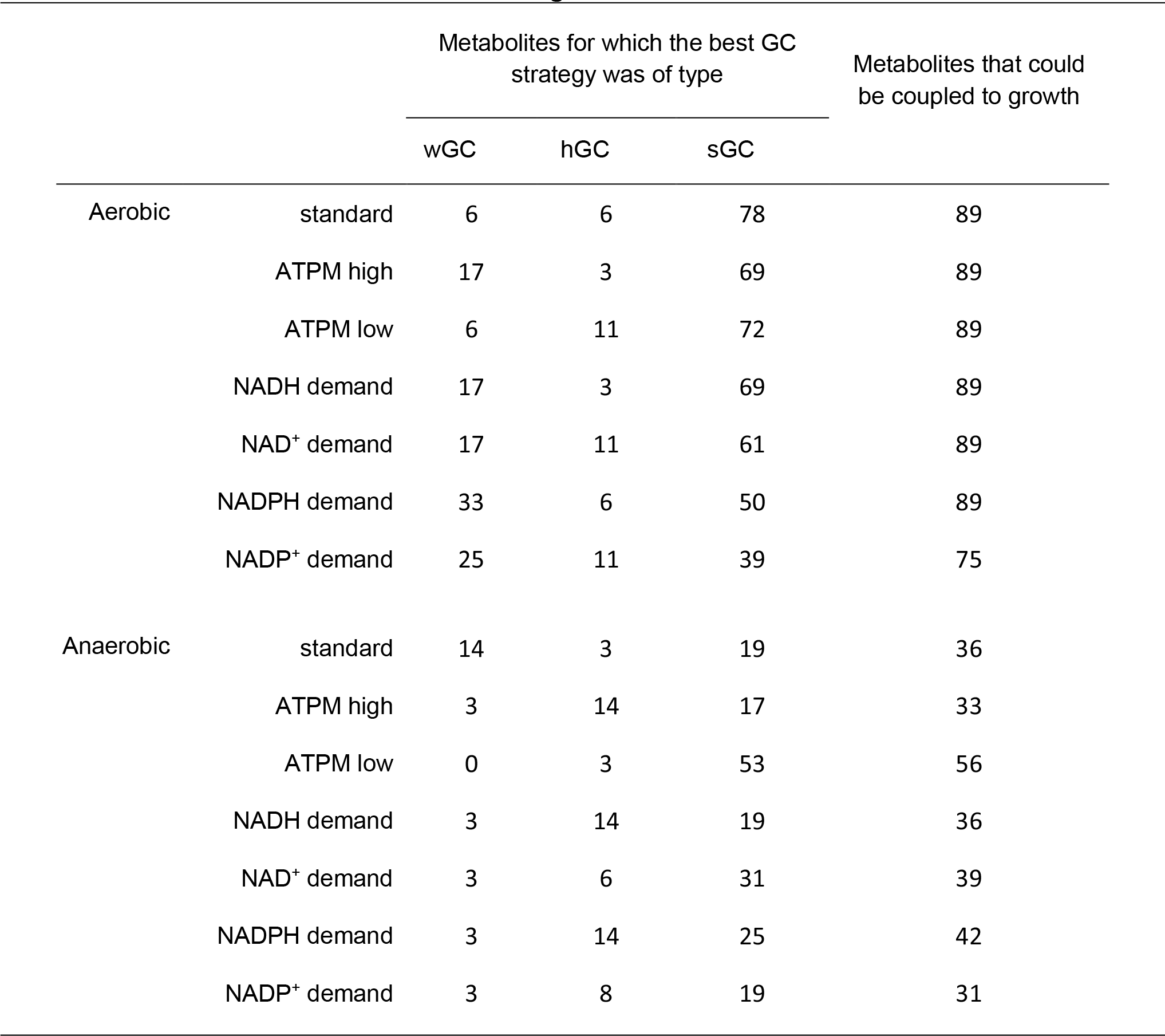
Relative numbers of metabolites for which strain designs of type wGC, hGC or sGC was identified. Strain designs were computed by gcOpt with the *E. coli i*AF1260 core model and glucose as the sole carbon and energy source. All values are given in percentage relative to the total number of investigated carbon metabolites of 36.

|  |  | Metabolites for which the best GC strategy was of type |  |  | Metabolites that could be coupled to growth |
| --- | --- | --- | --- | --- | --- |
|  |  | wGC | hGC | sGC |  |
| Aerobic | standard | 6 | 6 | 78 | 89 |
|  | ATPM high | 17 | 3 | 69 | 89 |
|  | ATPM low | 6 | 11 | 72 | 89 |
|  | NADH demand | 17 | 3 | 69 | 89 |
|  | NAD<sup>+</sup> demand | 17 | 11 | 61 | 89 |
|  | NADPH demand | 33 | 6 | 50 | 89 |
|  | NADP<sup>+</sup> demand | 25 | 11 | 39 | 75 |
| Anaerobic | standard | 14 | 3 | 19 | 36 |
|  | ATPM high | 3 | 14 | 17 | 33 |
|  | ATPM low | 0 | 3 | 53 | 56 |
|  | NADH demand | 3 | 14 | 19 | 36 |
|  | NAD<sup>+</sup> demand | 3 | 6 | 31 | 39 |
|  | NADPH demand | 3 | 14 | 25 | 42 |
|  | NADP<sup>+</sup> demand | 3 | 8 | 19 | 31 |

### Does product-coupled biomass precursor synthesis exhaust the GC potential?

A possible principle leading to GC, recently discussed by Jouhten *et al.* (2017), is the dependence of the synthesis of one or more biomass precursors on the activity of the target production, *e.g.*, by restricting precursor synthesis to reactions that split the substrate into a precursor essential for growth and the target metabolite. This assumption was tested by applying each found GC design to the iAF1260 core metabolic model and computing the capability of the impaired metabolic network to synthesize each reactant of the biomass synthesis equation while disabling the production of the respective target metabolite. In case the synthesis of a biomass precursor was blocked under these settings, the applied knockout strategy was considered to directly couple target compound production to precursor synthesis and thus to growth in general.

Sixteen biomass precursors were derived from the left-hand-side of the biomass formation reaction included in the *E. coli* core reconstruction. In Fig. 6, the percentage of accessible precursors for each identified strain design leading to GC is plotted against the GCS, not distinguishing between the number or reaction deletions or metabolites coupled to growth. For all strain designs showing a GCS below -1, thus being of type wGC, 100 % of the biomass precursors were still accessible. This contradicts the principle of a direct coupling between biomass precursor and product synthesis but is actually trivial since for wGC strategies product synthesis is only enforced above a certain threshold growth rate (cf. Fig 1). Likewise, this principle cannot explain product formation at zero growth for sGC. However, each identified sGC intervention strategy for anaerobic conditions resulted in blockage of *all* biomass precursors. Under aerobic conditions, this fraction was lower but still considerable. Only among the hGC strategies, a partial precursor blockage was found along with designs that had no effect on precursor availability at all. In none of the identified hGC solutions the synthesis of all biomass precursors was blocked.

**Figure 6.**
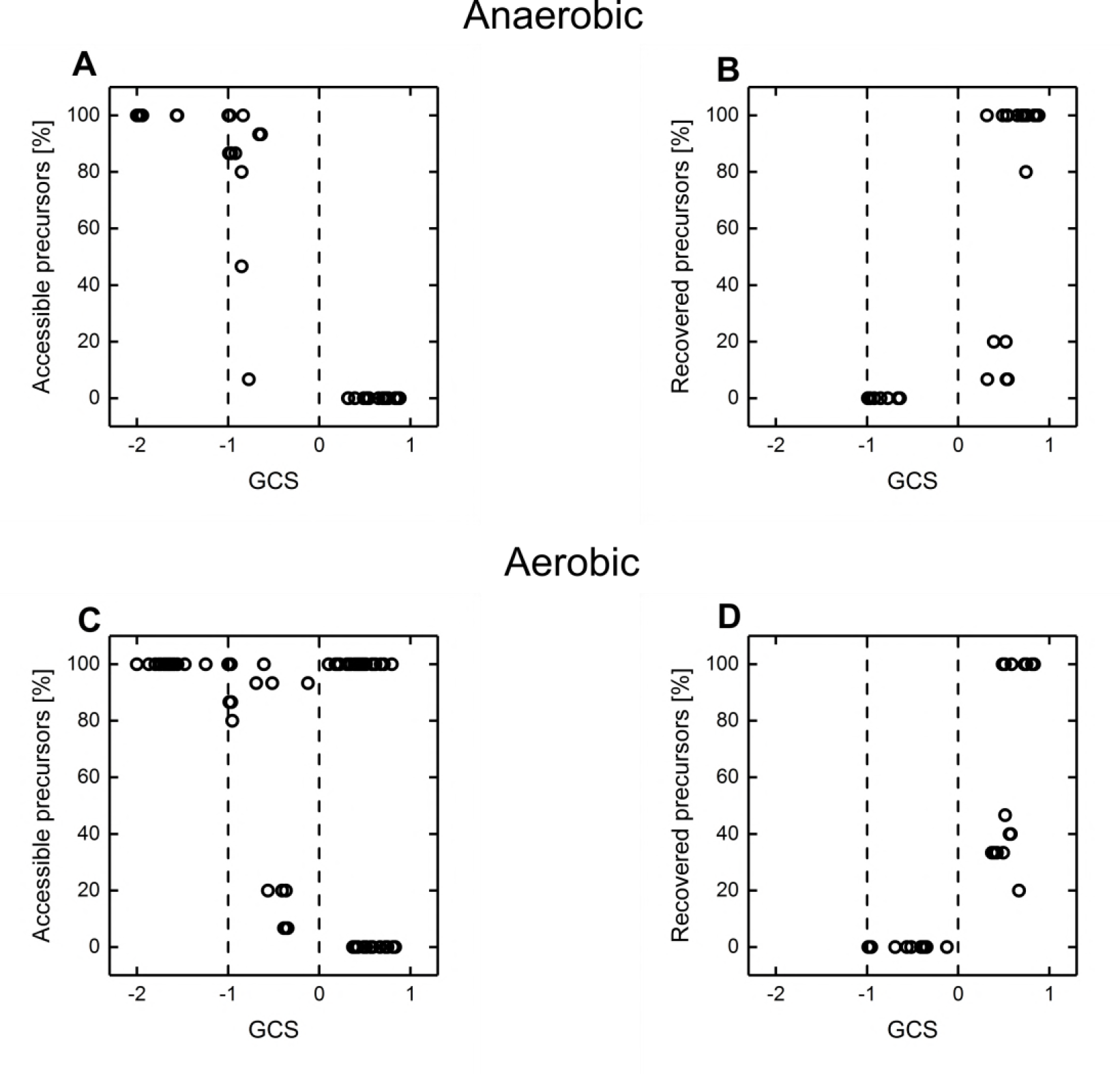
Biomass precursor availabilities for all identified GC strain designs under anaerobic and aerobic conditions for standard ATPM requirements and an unbounded, reversible ATP hydrolysis reaction. A Percentage of accessible biomass precursors for all identified GC strain designs of all metabolites and numbers of reaction deletions under anaerobe conditions as a function of the respective GCS. The vertical dashed lines separate the GCS range into three regions denoting wGC, hGC and sGC. B Cf. A, but for computing the biomass precursor availabilities a reversible, unbounded ATPM reaction was used allowing free generation and consumption of ATP. C Cf. A, but using the GC strains determined for aerobic conditions. D Cf. B, but using the GC strains determined for aerobic conditions.

Motivated by the observed increase in coverage and strength of GC strategies upon decreased ATP demand (Fig. 5), we wanted to further understand if and how the ATP metabolism might be a factor in establishing GC. To this end, we tested the biomass precursor availabilities for all identified strain designs allowing a reversible and completely unlimited flux through the ATPM reaction (Eq. 1). The consequence of this relaxation of the ATPM flux constraint is an unrestricted generation of ATP from ADP and free phosphate. While in all hGC cases (-1 < GCS < 0), none of the precursors could be recovered, *i.e.*, made accessible again by this relaxation, the synthesis of *every* blocked biomass precursor was restored for roughly 60 % and 80 % of the sGC strain designs (GCS > 0) under aerobic and anaerobic conditions, respectively (Fig. 6 B and D).

### The effects of relaxing cofactor balances on growth-coupling strain designs

The investigation of biomass precursor availability in the GC mutants indicated that an enforced production of the target compound (sGC) is likely due to a global metabolic necessity rather than caused by a strict dependence of the synthesis of a particular biomass precursor on target compound production. Moreover, ATP scarcity seemed to be a metabolic trigger for GC in those sGC cases in which the synthesis of any biomass precursor was blocked by the intervention strategies. To challenge this hypothesis, the GCS of a GC strain design was investigated upon relaxing the directionality constraint of the ATPM equation (cf. Eq. 1) thereby enabling the model to freely phosphorylate ADP to ATP and vice versa. Since the ATP metabolism is interconnected with the redox cofactor and cross-membrane proton balance, *e.g.*, via the electron transport chain and ATP synthase, a free NAD(P)H/NAD(P)^+^ generation and proton transport over the cell membrane were additionally tested for their effects on the GCS. To simulate this, the NADH and NADPH NGAM reactions (Eq. 2–3) were reintroduced and a new proton translocation reaction was added:

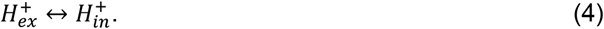

Here, the indices *ex* and *in* locate the H^+^ protons to the extracellular and intracellular compartment, respectively. Both reactions were unbounded, *i.e.*, allowed to carry any flux. All identified GC strategies and their GCS values under all investigated conditions are provided with Dataset EV1 and EV2.

For anaerobic conditions, GC was completely suppressed for all but two strategies by relaxing either the ATP balance, the NAD(P)H/NAD(P)+ conversion, the proton exchange or a combination of these strategies (Fig. 7). These two resistant strategies coupled formate to growth by forcing the carbon flux through the anchor reaction catalyzed by the pyruvate formate lyase, which splits pyruvate to formate and acetyl-CoA. However, the GCS of these strategies decreased when relaxing the constraints on cofactor generation and proton export.

**Figure 7.**
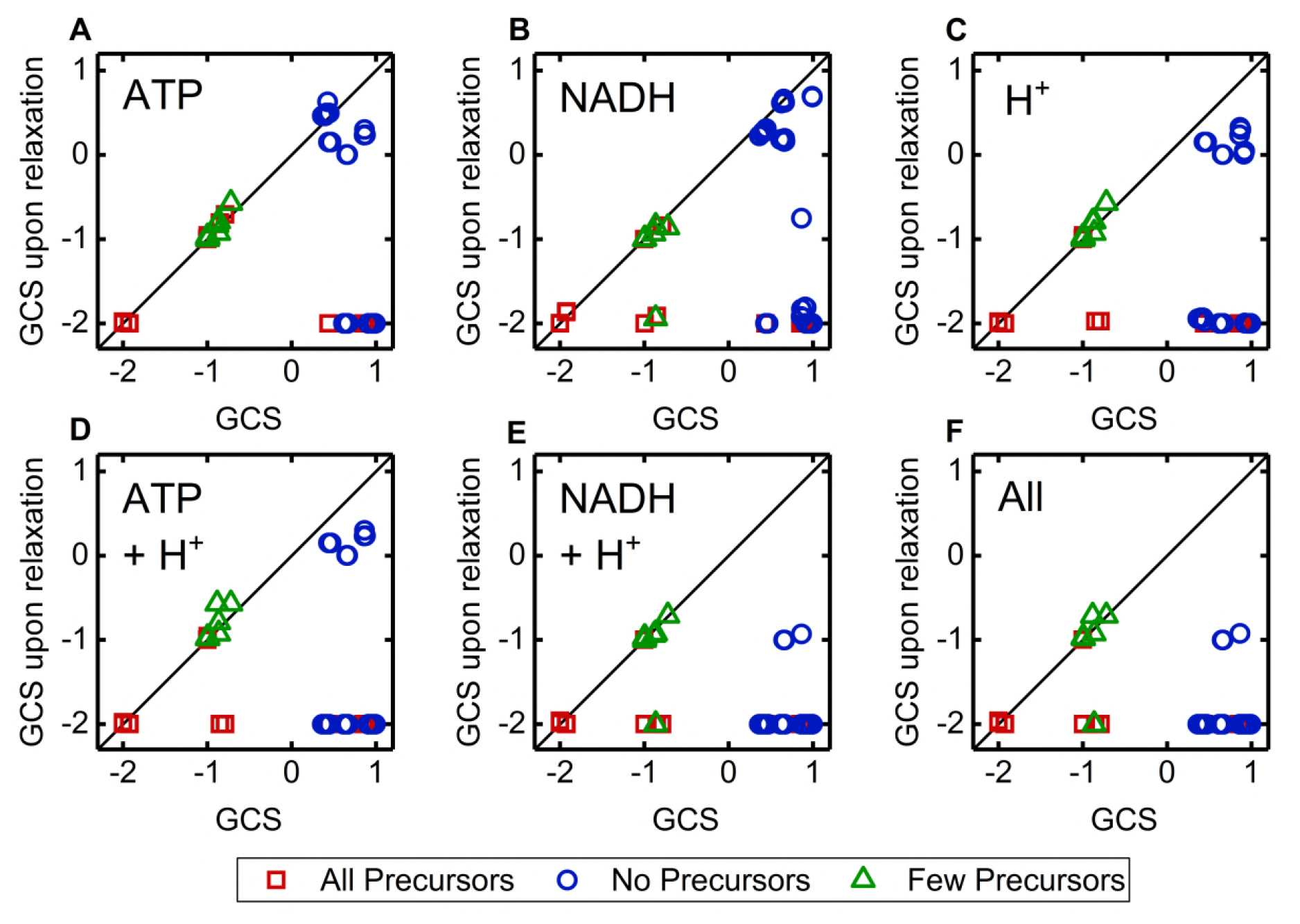
Comparison of the GCS of identified GC strain designs for anaerobic conditions and the corresponding GCS under relaxation of particular metabolic constraints (A-F). The color code relates to Fig. 6 showing the accessibility of biomass precursor for the same strain designs is shown. Red squares, blue circles and green triangles symbolize designs that allow for the synthesis of all, no or a reduced number of biomass precursors, respectively. A Relaxed ATPM constraint, enabling an unbounded de/phosphorylation of ATP/ADP according to Eq. 3. B Introduced Eq. 1 enables the re/generation of NAD/NADH. C H^+^ transport reaction allows for an unbounded translocation of H^+^ over the cell membrane. D Combination of A and C. E Combination of B and C F Combination of A, B and C.

Disclosure of the basic coupling principles was impeded by the interrelatedness of redox cofactor, ATP and H^+^ balancing (Fig. 9). For example, GC of lactate synthesis was abolished in most designs by relaxation of either ATP/ADP, NADH/NAD^+^ conversion or a free proton translocation. The basic coupling principle for this reduced metabolite is however NADH reoxidation, achieved by the reduction of pyruvate to lactate. Consequently growth-coupling is abolished upon opening the NADH balance. Relaxation of the ATP and proton balance had the same effect as it fuels flux through the NADH transhydrogenase, which couples NADH oxidation and NADPH reduction to proton import. The formed NADPH is oxidized in biomass forming reactions making NADH re-oxidation by lactate dehydrogenase activity superfluous. Under standard conditions NADH transhydrogenase activity is limited by the cell’s potential to maintain a proton gradient over the cell membrane. In contrast, GC of ethanol was only abolished when free proton exchange was enabled. That was not expected as ethanol and lactate share almost the same synthesis pathway and as ethanol is more strongly reduced than lactate. Apparently, GC of ethanol was achieved in these designs by coupling the intracellular proton balance to the ethanol-proton symporter activity. As all intervention GC strategies for ethanol included the deletion of the ATP synthase, proton export via a reversed ATP synthase activity under relaxed ATP turnover conditions was not possible. GC of pyruvate was diminished by both free proton transport and ATP/ADP conversion. Inspection of the flux distribution under relaxed ATP/ADP conversion conditions revealed that excess ATP was used to drive the ATP synthase as proton exporter. Consequently, pyruvate secretion was enforced by the need to balance intracellular protons as was the case for ethanol.

For aerobic conditions, relaxation of single or combinations of the tested constraints relieved GC for all wGC and most hGC strategies, as well. Again, formate was the only metabolite that was hard-coupled to growth by forcing flux through the pyruvate formate lyase anchor reaction. However, under aerobic conditions, this strategy is not of any relevance due to the pyruvate formate lyase’s sensitivity to oxygen (Edenharder, 1969). Surprisingly, more than 50% the sGC strategies were not affected by alleviating cofactor and proton supply although in most of these cases all biomass precursors were accessible without enforced product synthesis (Fig. 8). Inspection of the robust strategies showed that coupling of metabolites of the upper central carbon metabolism was achieved by prohibiting phosphoenolpyruvate (PEP) conversion in the lower central carbon metabolism by deletion of PEP carboxykinase and pyruvate kinase, and elimination of a cyclically operating pentose phosphate pathway, which would allow complete oxidation of the substrate to CO_2_. *In vivo*, this strategy might not be specific but could enforce the secretion of any upper central carbon metabolite. In our simulations, this was prohibited as only export reactions of lactate, ethanol and formate, were included in the model and the formation of these fermentation products was prevented by further reaction deletions in the GC strategies. In the residual designs the metabolism was curtailed in a way that forced glucose oxidation through metabolic anchor reactions, here, transketolase, transladolase or fructose bisphosphate aldolase, splitting the substrate into the target compound and an essential central carbon metabolism intermediate. As for the formate coupling strategies, the GCS of these strategies, although not completely abolished, was significantly reduced in most strategies upon relaxation of cofactor turnover and proton exchange. For the complete statistics corresponding to Fig. 7 and 8, we refer to Table EV2 and EV3.

**Figure 8.**
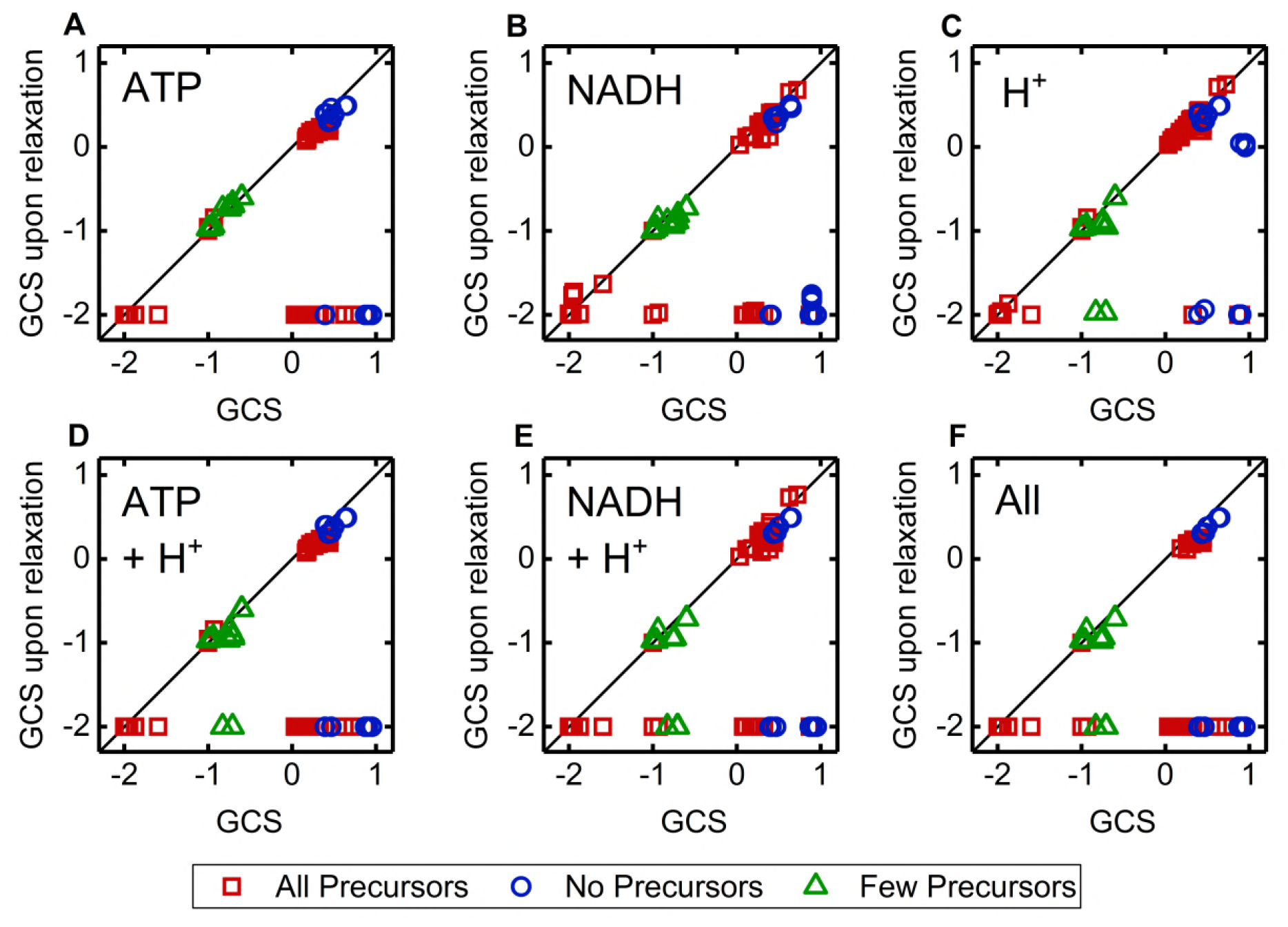
Comparison of the GCS of identified GC strain designs for aerobic conditions and the corresponding GCS under relaxation of particular metabolic constraints (A-F). The color code relates to Fig. 6 where the accessibility of biomass precursor for the same strain designs is shown. Red squares, blue circles and green triangles symbolizes designs that allow for the synthesis of all, no or a reduced number of biomass precursors, respectively. A Relaxed ATPM constraint, enabling an unbound de/phosphorylation of ATP/ADP according to Eq. 3. B Introduced Eq. 1 enables the re/generation of NAD/NADH. C H^+^ transport reaction allows for an unbounded translocation of H^+^ over the cell membrane. D Combination of A and C. E Combination of B and C F Combination of A, B and C.

### Growth-coupling affects the energy hierarchy of metabolites

The examination of the biomass precursor availability in the GC mutant strains and the influence of the ATP and NAD(P)H turnover and proton exchange on GC gave a first indication of the importance of the balancing of redox and energy cofactors. To further unravel the interdependency between GC and the cellular energy metabolism, the parameter ATP synthesis capability (ATPsc) was defined. ATPsc assesses the contribution of the product synthesis to the global provision or consumption of ATP (cf. Methods section and Fig. 9). The ATPsc represents the change in the maximal flux through the ATPM reaction when product synthesis is increased by 1 *mmol g*^−1^*h*^−1^. It is furthermore practical to normalize ATPsc with the number of carbon atoms of the target product yielding the ATPsc per carbon (ATPcsc). For the *E. coli i*JO1366 model the ATPcsc of CO_2_ is the highest followed by those of fermentation and overflow metabolites such as ethanol, lactate, succinate or acetate under anaerobic as well as aerobic conditions (Fig. EV2). This hierarchy therefore correctly reflects the order of metabolites secreted by this organism under oxygen-limited or carbon excess conditions (Clark, 1989, Varma et al, 1993). Hence, we used the energy hierarchy as a measure for quantifying the ATP gain from product synthesis relative to other possible side products.

**Figure 9.**
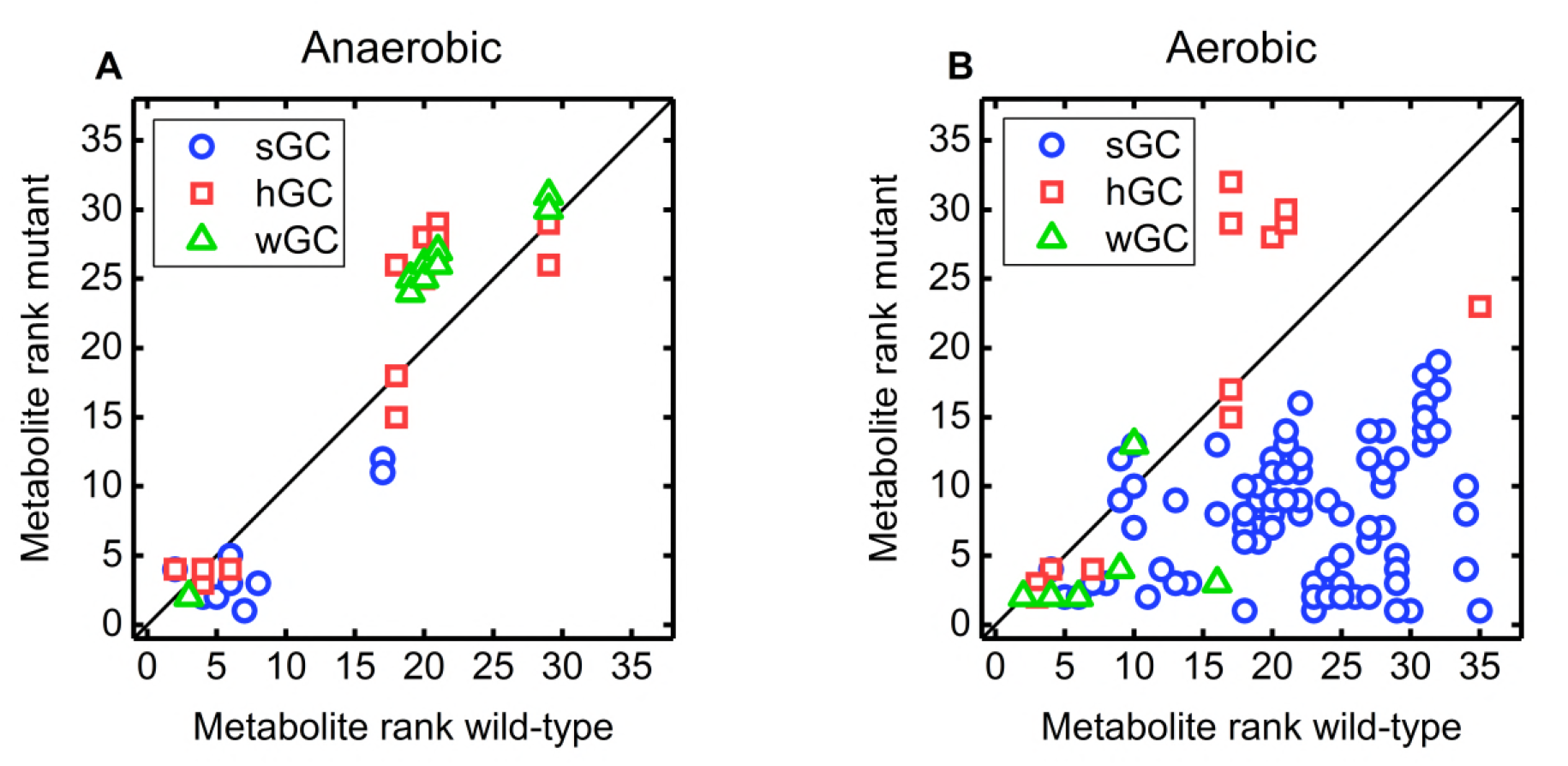
Comparison of the rank of GC target products of the *E. coli i*AF1260 core model in the energy hierarchy of metabolites of GC mutants and the wild-type. The hierarchy is based on the ATPcsc values. A GC strain designs calculated under anaerobic conditions. B GC strain designs calculated under aerobic conditions.

We hypothesized that growth coupling is induced or enhanced by the set of reaction deletions computed by gcOpt that deletes more efficient ATP generation routes so that the ATP yield of the target compound synthesis pathway excels those of residual ATP forming pathways. The observation that the ATP synthase was a frequent knockout target under aerobic conditions, diminishing ATP generation via oxidative phosphorylation, is in line with this hypothesis. Such a causality would become apparent by an increase of the ATPcsc in the mutant models and, in turn, an elevated rank of the target product in the energy hierarchy of metabolites. To test this assumption, the ATPcsc was calculated for a selection of 36 central metabolites of the wild-type *E. coli* iAF1260 core model and every identified GC intervention strategy using standard NGAM requirements. Energy hierarchies of metabolites for each GC strain design were arranged such that the metabolite with the highest ATPcsc value was ranked position one.

Under anaerobic and aerobic conditions, for most sGC strategies the target compound’s rank in the hierarchy of metabolites increased compared with wild type conditions (Fig. 10). This was reversed for the wGC and hGC designs, in which the majority of target products faced a reduction of the energy hierarchy rank. For aerobic conditions (Fig. 9 B), the upward shift of the target compounds in the energy hierarchy was considerably more pronounced than under anaerobic conditions and for 30 % of the sGC strain designs the target compound was even ranked second or first, thus, in case of the latter, surpassing CO_2_ as the most beneficial production pathway for generating an excess supply of ATP.

## Discussion

Two major GC principles have been described in the literature, in which target production is enforced by either coupling it to the synthesis of biomass precursors or the balancing of energy and redox cofactors. Besides, the observation that yield and specific productivity is increased upon inducing an ATP futile cycle was discussed as another principle (Hädicke *et al*, 2015; Ebert *et al,* 2011). However, this latter interrelationship cannot be transferred to GC according to the findings shown in this work. Under anaerobic conditions, only a simulated *decrease* of the ATP consumption for cell maintenance processes led to an increased number of growth-coupled metabolites as the surplus of available ATP allowed for the secretion of compounds with higher energy contents such as phosphorylated intermediates. Identified GC strain designs also showed enhanced GCS at reduced maintenance requirements. The prevalent ATP scarcity under anaerobic conditions may also be the reason for the observed stagnation of the GCS with increasing size of the intervention sets. A metabolic boundary, most likely ATP limitation, might confine the maximal ratio between the energetically disadvantageous product synthesis and growth. ATP supply is less critical in an aerobic setup due to a fully functional respiratory chain. This is reflected in the indifference of the GCS to alterations of energy and equally redox cofactor. Apparently, the better supply of redox and energy cofactors under aerobic conditions implies a superior metabolic capacity for product-growth coupling as was previously hypothesized (von Kamp & Klamt, 2017). Accordingly, we can phrase the following GC principle:

**<u>Principle 1:</u>** *Generation of a strong coupling between growth and product synthesis is ultimately limited by the cell’s natural capacity to regenerate energy equivalents in form of ATP.*

This became also apparent from the ATPcsc values of GC mutants. For aerobic conditions, most target products soared in the energy hierarchy ranking. Thus, their synthesis pathway became a more, or in some case *the* most advantageous metabolic route to regenerate ATP in terms of an optimal compromise between carbon usage and ATP yield.

In summary, the here presented, rigorous calculations of GC strategies using gcOpt confirmed previously published results (Klamt & Mahadevan, 2015; von Kamp & Klamt, 2017):

**<u>Principle 2:</u>** *Feasibility of GC holds for a wide range of metabolites.*

Yet, GC of energy-rich as well as oxidized metabolites and the ability to reach high coupling strengths was found to be limited to aerobic conditions.

The apparent, global feasibility of GC pronounces the applicability of this concept for *any* microbial strain engineering project aiming to increase productivity and yield. The intuitive approach is to enforce an obligatory dependence of the synthesis of one or more essential biomass precursors on target compound production. However, such a GC criterion can only explain or lead to hGC characteristics, in which product synthesis is strictly bound to biomass formation. In fact, we found that the majority of hGC strategies blocked the synthesis of one or more biomass precursors at zero product formation. The concept can be broadened to the following principle and was indeed evident for 50% of all aerobic hGC and sGC strategies.

**<u>Principle 3:</u>** *Linking product synthesis to reactions essential for any steady state flux distribution on the chosen carbon source results in holistic and strong GC. Those reactions include but are not restricted to biomass precursor forming reactions.*

However, our analysis also highlights the balancing of global cofactor as an additional, important criterion for establishing or enhancing GC, hence leads us to a specialized version of the third principle.

**<u>Principle 4:</u>** *Reconfiguration of the metabolic network rendering the product synthesis pathway the superior ATP supply route is one major principle for generating strong growth coupling or enhancing the GCS.*

This can be intuitively inferred from the observation that the ATP synthase is a frequent knockout target in aerobic GC strategies and more quantitatively be described with the rise of the target metabolite in the energy hierarchy. This is supported by the observation of Jensen & Michelsen (1992) that an ATP synthase deficient *E. coli* strain shifts the flux distribution towards substrate-level phosphorylation pathways, *i.e.* glycolysis and TCA cycle, and the secretion of correlated metabolites. We conclude that ATP synthase deletion forms a basis for GC under aerobic conditions whereas additional knockouts enforce specificity of product secretion as can be derived from the steadily increasing mean GCS with increasing genetic interventions. Exceptions from this pattern are fermentation products or products exported via proton symporters, for which GC is induced by disrupting alternative NADH re-oxidizing or proton translocating pathways.

Concludingly, metabolic network reconfigurations that render product secretion into a carbon drain necessary for metabolic activity might be the more robust GC approach as it is independent of cofactor and proton balancing that might vary under different growth conditions. However, such a coupling might not be possible for all metabolites. Our analysis revealed that coupling product formation to cofactor supply or turnover not only enhances the GCS of the former strategies but also seems to be globally applicable to any metabolite. Such metabolic designs that are based on such cofactor balancing are hard to derive manually due to the complex interconnectedness of energy and redox cofactors within metabolic networks. Accordingly, we argue that computer-aided network analysis can accelerate the development of strain designs strictly coupling production to microbial growth by predicting effective GC strategies with a reasonable number of gene deletions.

## Methods

### Formulation of gcOpt

gcOpt is geared to existing multi-level optimization frameworks and their lower-level formulations optimizing an engineering objective by searching for appropriate sets of reaction deletions (Burgard *et al,* 2003; Tepper & Shlomi, 2009). gcOpt maximizes the minimally guaranteed production rate *v_t_* of a target compound *t* for a fixed growth rate *µ*_*fix*_. The corresponding bi-level optimization problem is formulated as follows:

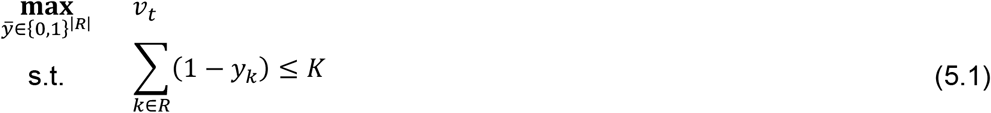

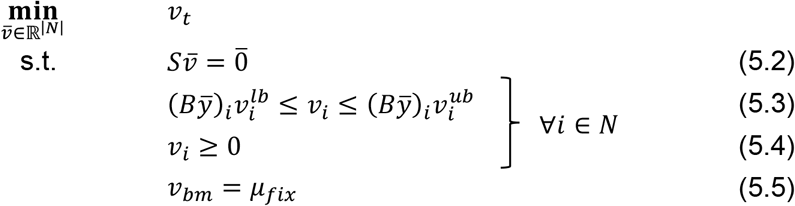

where *ӯ* is a boolean vector indicating for each reaction *i* ∈ *R* within the reversible metabolic model if *i* is inactive (0) or active (1). With constraint (5.1), the size of the reaction deletion set is limited to *K* interventions. Note that the gcOpt formulation requires the splitting of all reversible reaction of the original model into irreversible forward and backward reactions. This results in an irreversible metabolic model containing *N* reactions with strictly positive fluxes (Eq. 5.4). The flux value of each irreversible reaction is contained in the vector 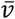. Steady state mass balances of intracellular metabolites are assured by Eq. 5.2. Here, 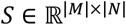 is the stoichiometric matrix where each non-zero value *S_j,i_* denotes the stoichiometric coefficient of metabolite *j* ∈ *M* participating in reaction *i ∈ N.* Each flux *v_i_* is constrained by lower and upper bounds 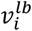 and 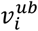, respectively, or set to zero if *y_k_* indicates a knockout (Eq. 5.3). The connection between a reversible reaction *k* and one of its irreversible counterparts *i* is kept in the mapping matrix *B* ∈ {0,1}^|N|×|R|^ by *B_i,k_* = 1. Since the biomass reaction *v_bm_* is fixed to *µ_fix_* by Eq. 5.5, *µ_fix_* needs to be lower than the maximally achievable biomass formation rate 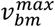.

Solving the nested mixed-integer optimization problem (Eq. 5) using linear programming solvers is intractable due to the high number of reactions, the fluxes of which being the variables in (1) (Burgard *et al,* 2003). By virtue of the linearity of the outer and inner objective function as well as the posed equality and inequality constraints, the bi-level optimization problem (Eq. 5) can be recast to a single-level mixed integer linear program (MILP) by exploiting the strong duality theorem in linear programming (Burgard & Maranas, 2003). For gcOpt, the reformulation was done as described by Tepper & Shlomi (2009) but adapted to the differently formulated objective functions.

In this work, core models of the central carbon metabolism introduced by Trinh *et al.* (2008) and Orth *et al.* (2010) as well as the advanced metabolic model iJO1366 of *E. coli* K-12 MG1655 (Orth *et al,* 2011) were used. To improve the tractability of strain designs computations, the solution space of the genome-scale model iJO1366 (1366 genes, 2251 reactions and 1136 metabolites) was reduced following a preprocessing routine similar to a protocol of Feist *et al.* (2010), which was integrated into the gcOpt framework. More specifically, essential as well as exchange, diffusion, transport and spontaneous reactions were excluded from the set of possible target reactions for deletion. Furthermore, reactions contained in the subsystems cell envelope biosynthesis, membrane lipid metabolism, murein biosynthesis, tRNA charging and glycerophospholipid metabolism were also not regarded as deletion targets. In addition to reducing the solution space of the problem posed by gcOpt, the actual number of reactions within the considered metabolic model was trimmed by entirely removing all reactions unable to carry any flux. These so called blocked reactions were identified by flux variability analysis (Mahadevan & Schilling, 2003) by means of a maximum and minimum flux equal to zero.

The gcOpt framework was implemented in Matlab (R2016b, The Mathworks, Newark) and is freely available on GitHub (Alter, 2018). For solving the single-level MILP derived from problem (1), the Gurobi Optimizer (7.0.2, Gurobi Optimization, Inc.) was utilized. All computations in this work were conducted on a Windows 7 machine with a maximum configuration of 16 GB of RAM and a AMD FX-8350 Eight-Core (à 4.00 GHz) processor.

### Quantification of the growth-coupling strength

To quantify and compare the GC level or strength of microbial strain designs, GCS, a novel measure for the growth coupling strength based on the yield space representation, was defined. As visualized in Fig. 4, the ratio between the area *IA* below the lower yield bound and the total area *TA* under the upper yield hull curve in the yield space of a strain design determines the GCS. This expresses the principle that the zero- or low-yield flux modes are made inaccessible the stronger the coupling between growth and product synthesis. In addition, the minimally guaranteed target product yield at maximal growth 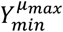 divided by the theoretical maximal yield *Y_max_* is considered as a factor in the formula for the GCS (Eq. 6.1 – 6.3). For strain designs with similar GC levels according to an evaluation of the yield space areas, this factor promotes those that guarantee high yields at elevated growth rates. To be able to directly distinguish the GC types sGC, hGC and wGC, the intersection of the lower yield bound and the growth axis was further integrated. Following this, the GCS is finally calculated as follows:

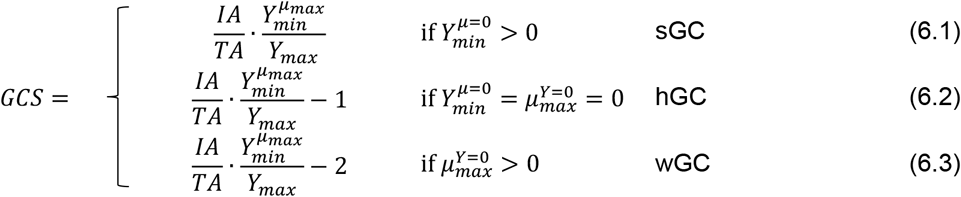

Here, 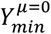 and 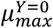 are the minimal target product yield at zero growth and the maximal growth rate at zero yield, respectively. GCS increases with increasing GCS values and the three GC types are defined by distinct GCS ranges. To allow for an immediate distinction of the GC types, GCS of hGC and wGC strategies are normed by considering one and two as an additional subtrahend in Eq. 6.2 and 6.3, respectively. Thus, a GCS between -2 and -1 denotes wGC, the interval [–1,0] indicates hGC and GCS > 0 implies sGC. Hence, the GCS parameter enables both a qualitative classification and a quantitative ranking of GC strain designs. In the data evaluation process, strategies with GCS ≤ -1.975 and between -0.975 and -1 were considered to confer no coupling.

### Probing the biomass precursor availability

To evaluate the capability of a metabolic network to synthesize a particular biomass precursor the biomass synthesis equation was singularized into separate, independent reactions, *i.e*., for each single reactant *M* with stoichiometric coefficient *v* in the biomass equation a new, unbounded reaction of the form *vM* → was defined. Similarly, for each product *N* and stoichiometric coefficient *w* a reaction → *wN* was added to the model. If the consumption of a precursor was coupled to the production of a certain compound, *e.g.* ATP and ADP, both were linked in a new reaction of the form *vM* → *wN.* The original biomass equation was erased from the model. The availability of a biomass precursor of this original equation was then tested by setting up a linear program that maximized the related singularized reaction subject to all mass balance, substrate uptake and thermodynamic constraints of the original model. A maximal achievable flux of zero, implies loss of the metabolic capacity to synthesize the respective precursor and hence, this precursor is inaccessible. For cases with positive maximal fluxes, the synthesis of the precursor is not impaired.

### ATP synthesis capability

The ATP synthesis capability (ATPsc) parameter was created to deduce the influence of byproduct secretion on the synthesis and provision of the cellular energy equivalent. For a given metabolic network and production rate of a target compound, the ATPsc describes the change of the maximal flux through the reaction ATPM in response to a change of the production rate of the target chemical. Mathematically, the ATPsc is defined by the derivative *dv_ATPM max_/dv_target exchange_*, with *v_ATPM max_* being the maximal flux through the ATPM reaction (Eq. 1) computed by linear programming and a metabolic network constrained with a target product exchange rate fixed to values between zero and the maximal flux. Graphically, ATPsc can be determined by plotting the ATPM flux values against the product exchange rate and calculating the slope of the graph (Fig. EV1). Using the *E. coli* iJO1366 metabolic model, the ATPsc was calculated for a range of metabolites of the central carbon metabolism for low production rates. Thus, the resulting ATPsc values correspond to the differences in maximal accessible excess ATP between inactive and active metabolite secretion. For each metabolite, the ATPsc was calculated for a range of accessible growth rates. Fig. EV2 shows the results for anaerobic and aerobic conditions. Here, mean ATPsc values were normalized by the number of carbon atoms of the target compound (termed ATPcsc).

## Acknowledgments

We thank Lars Küpfer for critically reading and commenting the manuscript and helpful discussion.

## Author contributions

TBA and BEE conceived and designed the study and evaluated the simulation results. TBA developed the gcOpt algorithm and performed all simulations. TBA and BEE wrote the manuscript. LMB supervised the study and edited the manuscript. All authors read and approved the final paper.

## Conflict of Interest

The authors declare that they have no conflict of interest.

---

**Figure EV1.**
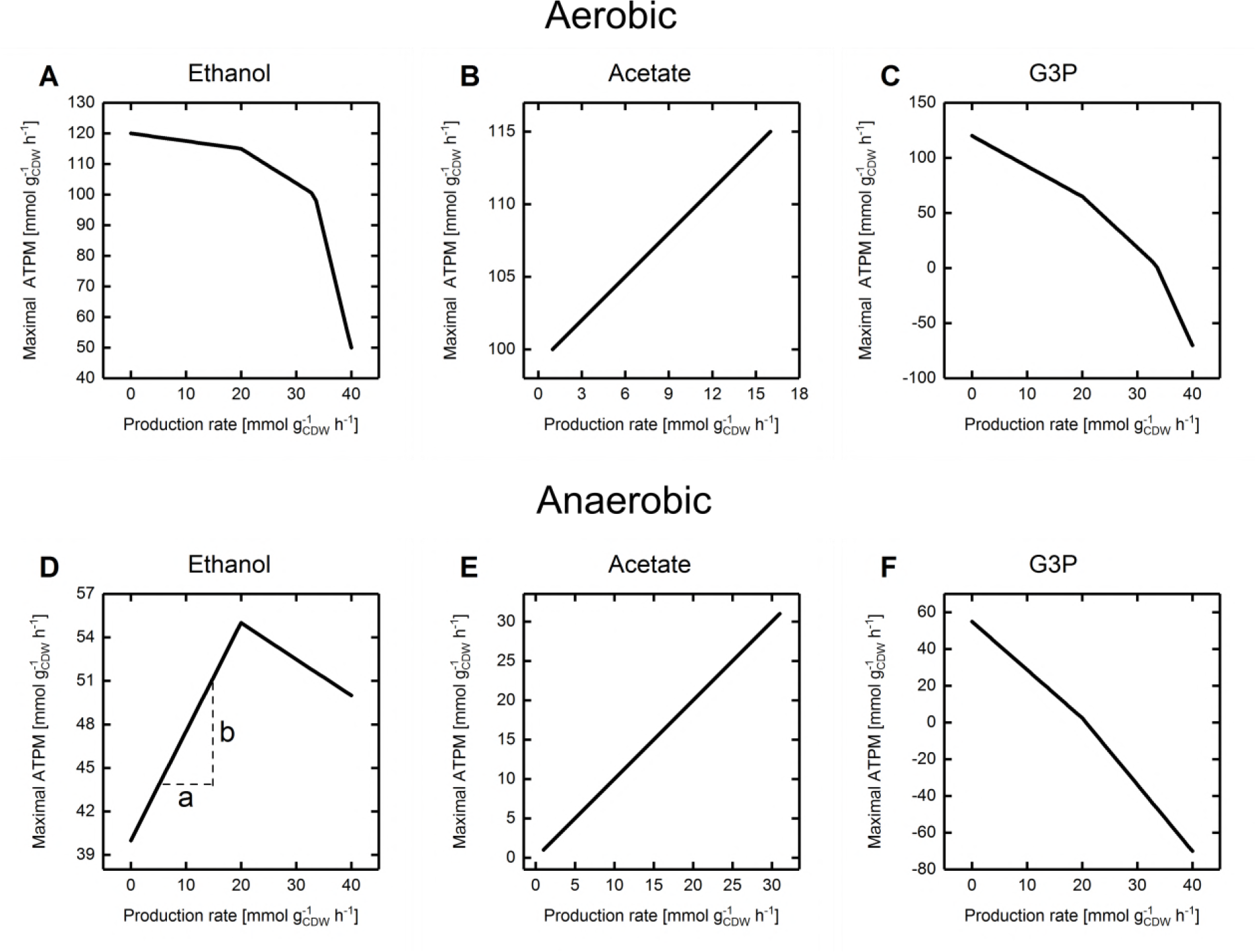
Relation between maximal ATPM and the production rate of several metabolites under anaerobe and aerobe conditions using the *E. coli* /AF1260 core model and glucose as the sole carbon substrate. A-C Anaerobic condition D-F Aerobic condition. In D, calculation of the gradient *s* is shown exemplarily by *s* = *b*/*a*.

**Figure EV2.**
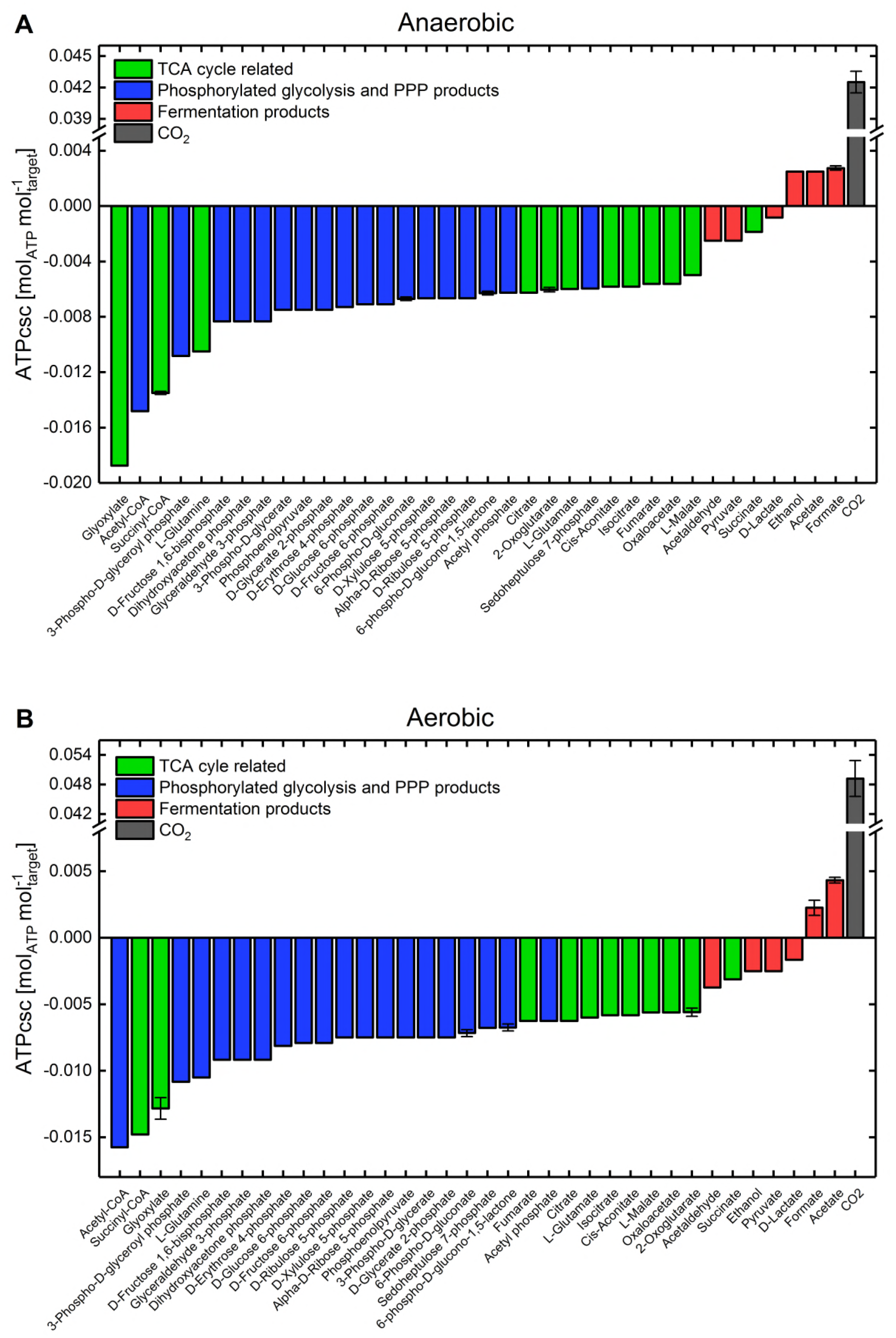
ATPcsc values for several metabolites of the central carbon metabolism using the *E. coli i*JO1366 metabolic model and glucose as the sole carbon and energy substrate. Error bars denote the standard deviation of ATPcsc calculations at different growth rates spanning the feasible range of growth states. The color code links the metabolites to glycolysis and pentose phosphate pathway (PPP) (blue), TCA cycle (green) and fermentative pathways (red), respectively. A Anaerobic condition B Aerobic condition

